# Increased conductance associated with higher photosynthesis in leaves and green stems of avocado exposed to diffuse light

**DOI:** 10.1101/2022.09.09.507344

**Authors:** Z. Carter Berry, Eleinis Ávila-Lovera, Kendra Ellertson, Gregory R. Goldsmith

**Author notes:** Contact: Z. Carter Berry, Wake Forest University, Winston Hall, 1834 Wake Forest Road, Winston-Salem, NC 27109.

## Abstract

Research has demonstrated that diffuse light drives changes in leaf photosynthesis, with the direction and magnitude varying across species; however, our understanding of the relationship between diffuse light and plant gas exchange, as well as the mechanisms driving these relationships remain unresolved. We studied the effects of diffuse light on plant function in potted individuals of *Persea americana* (avocado). We first measured leaf gas exchange subject to varying proportions of direct and diffuse light, as well as photosynthetic response to varying CO_2_ (A-C_i_ curves) in predominantly direct and predominantly diffuse light. We find that leaf photosynthetic rates increase as the proportion of diffuse light increases and that those changes are associated with stomatal conductance, rather than photosynthetic biochemistry. Given that avocados have green stems, we then measured stem gas exchange in predominantly direct compared to predominantly diffuse light. While we also observed an increase in conductance in stems subject to diffuse light, there was not an increase in photosynthetic rate, effectively decoupling gas flux from carbon gain. Finally, by scaling measurements of gas exchange to the plant, we demonstrate that stem bark conductance contributes proportionally more to whole-plant conductance under diffuse light. Our results add to our understanding of the potential mechanisms that govern how plant function varies in response to changes in light quality, the first paper to demonstrate mechanisms to explain increases under diffuse light. As diffuse light increases globally, this variable needs to be integrated into our understanding of plant carbon-water tradeoffs in response to climate change.

## Introduction

For plants, not all light is equal. In addition to quantity and spectral quality, we now understand that the angle of light arriving at the plant leaf or stem surface (i.e., diffuse light independent of quantity or spectral quality) can significantly alter plant gas exchange and that the directionality and magnitude of this effect varies among species (Earles et al. 2017, Berry & Goldsmith 2020, Berry et al. 2022). However, the relationship between plant gas exchange and varying proportions of direct-to-diffuse light, as well as the mechanisms governing differences in plant gas exchange in diffuse light, remain relatively unexplored.

Determining the relationship between plant gas exchange and varying proportions of direct/diffuse light can be pursued by both observational and experimental approaches. The primary evidence comes from observations of net ecosystem exchange under varying sky conditions (Roderick et al. 2001, Gu et al. 2002, Misson et al. 2005, Alton et al. 2007, Mercado et al. 2009, Williams et al. 2014, Cheng et al. 2015, Baguskas et al. 2021). These studies demonstrate increased productivity in diffuse light conditions, but with an inability to tease apart the driving mechanisms. A recent analysis found that simply incorporating the diffuse fraction of light into a land surface model improved predictions of primary productivity by 50% (O’Sullivan et al. 2021). While fewer in number, most experimental approaches have measured leaf gas exchange after exposing leaves to a predominantly direct or predominantly diffuse light source Brodersen & Vogelmann, 2007; Brodersen & Vogelmann, 2010; Earles et al., 2017). Only recently has the ability to experimentally measure the effects of intermediate levels of diffuse light on plant gas exchange become available (Berry et al. 2022).

Several different mechanisms for changes in leaf gas exchange in diffuse light conditions have been proposed. Early research hypothesized that diffuse light would limit photosynthesis by decreasing the transmittance of light through the leaf (Brodersen & Vogelmann, 2007; Brodersen & Vogelmann, 2010; Earles et al., 2017). While limited light transmittance may play a role, it cannot be the only potential explanation, particularly given that some species have been observed to increase photosynthetic rates under diffuse light (Berry & Goldsmith 2020, Berry et al. 2022). A second hypothesis for changes to photosynthesis in diffuse light is that it is driven by biochemical changes to photosynthetic efficiency through greater electron transport rates (*J*_*max*_) or the maximum rates of carboxylation (*V*_*cmax*_) (Oguchi et al., 2011, Hogewoning et al., 2012; Earles et al., 2017). Finally, a third hypothesis that could explain changes to photosynthesis is changes to stomatal conductance. Stomatal conductance, as governed by guard cells, can change quickly in response to intensity and wavelength of light, particularly blue (and to a lesser extent red) light (Ogawa 1981, Assmann 1988, Shimazaki et al., 2007). A recent review clearly laid out these hypotheses, but they have not been explored empirically (Durand et al. 2021).

Most research on diffuse light has focused on leaves, however, many woody plant species can also have green stems (Nilsen 1995, Ávila et al. 2014). Having green stems provides physiological advantages: green stems represent an additional source of carbon, they decrease the amount of CO_2_ being released to the atmosphere, and, in many species, they are more water-use efficient than leaves (Ehleringer et al. 1987, Nilsen and Sharifi 1997, Ávila-Lovera and Tezara 2018, Ávila-Lovera et al. 2018). Understanding diffuse light responses in green stems is particularly relevant, as most stems are surrounded by leaves and are thought to receive diffuse light and therefore be “shade-acclimated” (Cernusak and Marshall 2000, Aschan et al. 2001, Wittmann et al. 2001, Damesin 2003, Yiotis et al. 2008). Further, light transmittance is considered a major limiting factor for photosynthesis in stems, which have additional layers to pass through to reach chloroplasts (Sun et al., 2003; Teskey et al., 2008; Ávila et al., 2014).

In this study, we seek to understand how (1) photosynthesis varies with differing proportions of diffuse light in leaves and stems, (2) begin to explore the mechanisms that might explain variation in photosynthetic rates under diffuse or direct light, and (3) quantify the magnitude of these effects at the scale of the whole plant. To do this, we quantified the photosynthetic response to diffuse light, the photosynthetic efficiency (*V*_*cmax*_ and *J*_*max*_), and conductance to water vapor in leaves (stomatal) and stems of common avocado (*Persea americana* Mill.). We predicted that photosynthesis would decrease with increasing proportions of diffuse light because of reduced transmittance in this species with thick leaves (Earles et al. 2017). We also expected this reduced light transmittance through the leaf in diffuse light would reduce the maximum rate of Rubisco carboxylation (*V*_*cmax*_) and the maximum rate of electron transport (*J*_*max*_). In contrast, we predicted that stems might exbibit higher photosynthesis with diffuse light since they may be acclimated to such conditions.

## Methods

### Measurement Design and Species

The experiment was conducted on 12 potted *Persea americana* (avocado) trees on the campus of Chapman University in Orange, California. We chose avocado because our previous work on diffuse light focused on tropical species (Berry and Goldsmith 2020), because avocados have green stems that are known to carry out stem CO_2_ reassimilation (Valverdi et al. *in review*), and because of the high commercial value of the species. The trees were obtained in May 2020 from Maddock Ranch nursery in Fallbrook, CA, and were 1 year old. All measurements were made between June and September 2020. All trees were between 1.7 and 2 m tall with a basal stem diameter ranging from 39 to 48 mm. All plants were in 15-gallon pots and were watered daily to field capacity.

### Photosynthesis and respiration measurements

To quantify the effects of diffuse light on rates of leaf and stem carbon assimilation, instantaneous gas exchange measurements were made on leaves and stems experiencing direct and diffuse light conditions. We used a portable infrared gas analyzer (Li-Cor 6800; Li-Cor Inc. Lincoln, NE, USA) with a modified integrating sphere attached to the leaf chamber to create diffuse light environments. The integrating sphere mounts directly on top of the leaf chamber and allows for the light source to be mounted at five different positions from directly above the leaf (predominantly direct light) to perpendicular to the leaf (predominantly diffuse light). The interior of the sphere has a highly reflective coating to scatter light that reflects off the surfaces. By moving the light along these positions, we were able to create light conditions that maintained the same intensity, but varyied the proportion of direct light from 77% to 29% (for more information, see Berry et al. 2022).

Instantaneous gas exchange measurements were made on one fully expanded, mature, and healthy leaf between 08:00 and 13:00. Leaves were placed in the 6 × 6 cm large leaf chamber (Model 6800-13; Li-Cor Inc. Lincoln, NE, USA) with the accompanying red (65%), green (10 %), blue (20%) and white (5%) LED light source and allowed to acclimate at a photosynthetically active radiation (PAR) of 1295 μmol m^-2^ s^-1^ until photosynthesis was stable. The chamber was set at a temperature of 27 °C, CO_2_ of 410 μmol mol^−1^, relative humidity of 50%, fan speed of 10,000 rpm, and a flow rate ranging from 500 to 1000 μmol s^−1^. Once the measurement was recorded, the light source was moved to the next position on the sphere and again allowed to stabilize for at least 5 minutes. This process was repeated at all five light positions for all 12 individuals.

For stems, instantaneous gas exchange rates were only measured in predominantly direct and predominantly diffuse light in 1-2 stems from each of the 12 plants. Because avocado does not have stomata on its green stems, the carbon flux represents net CO_2_ exchange considering photosynthesis and respiration. By subtracting light respiration from dark respiration, we estimated a CO_2_ re-assimilation rate (Cernusak and Marshall 2000). Measurements were done on stems between 4 and 6 mm thick with an internode length > 6 cm to be able to accommodate the 6 × 6 cm leaf chamber. A combination of foam tape and terostat butyl tape was used to seal the chamber around the stem allowing for accurate measurements of gas exchange in intact stems, an improved protocol from Ávila-Lovera et al. (2017). Chamber conditions were the same as for leaves, except for flow rates, which were set to 300 μmol s^−1^ to accommodate low rates of gas diffusion in stems without stomata. For all three measurements (direct, diffuse, and dark), values of net CO_2_ exchange rate and stem bark conductance were allowed to stabilize for at least 20 min before logging the values.

### A-C_i_ curves

To quantify the effects of diffuse light on leaf photosynthetic biochemistry, A-C_i_ curves were constructed using the rapid A-C_i_ response (RACiR) method (Stinziano et al. 2017, Lawrence et al. 2018, Saathoff et al. 2021). This non-steady-state method differs from traditional A-C_i_ curves in that it continually changes the CO_2_ concentration in the chamber in a stepwise fashion. To account for offsets between gas analyzer measurements and CO_2_ concentrations in the chamber under this continually changing condition, an identical A-C_i_ curve is run with an empty chamber. From this empty chamber dataset, a polynomial is fit that is used as a correction factor for the data with the leaf in the chamber. We measured RACiR A-C_i_ curves ranging from CO_2_ set between 10 to 800 μmol mol^−1^ over 9 min, corresponding to a linear increase in CO_2_ of 87.8 μmol mol^−1^ min^-1^. The calculation of C_i_ based on these parameters produced A-C_i_ curves that ranged from C_i_ of 55 to 550 μmol mol^−1^. Data for all curves were logged every 5 sec. Chamber conditions were set to a leaf temperature of 27° C, relative humidity of 50%, and light level of 1295 μmol m^-2^ s^-1^. Leaves were allowed to acclimate at a reference CO_2_ of 410 ppm prior to beginning each RACiR curve. All parameters were kept identical for the runs with leaves and empty chambers. One RACiR curve was run for each leaf with predominantly direct light and one curve in predominantly diffuse light on each of the 12 trees.

### Estimating the role of leaves and green stems to whole-plant carbon flux and conductance

To quantify the relative contribution of leaves and stems to total carbon fluxes and conductance to water vapor in direct and diffuse light conditions, we calculated the total photosynthetic area of leaves and stems for each plant (Ávila-Lovera et al. 2020). One large branch was harvested from each plant and the total length, diameter at the base, and the diameter at the tip was measured. From these measurements, we were able to estimate the surface area of each stem approximated as a cone. We then counted the total number of branches to determine total branch surface area. We used the same approach for the main stem of the tree by measuring the total height, diameter at the base, and the diameter at the tip. For leaves, we calculated the area of 4 leaves per branch and counted the total number of leaves to determine a total leaf area of the plant. Using these areas, we then calculated the total contribution of leaves and stems to the conductance to water vapor under direct and diffuse light.

### Data Analysis

Relationships between the angle of diffuse light and gas exchange parameters of leaves were all fit with linear models. The A-C_i_ curves were fit using non-linear regression. To compare differences in diffuse and direct light gas exchange and A-C_i_ parameters, we conducted a two-tailed paired-samples t-test. This included comparisons of net CO_2_ exchange rate and conductance to water vapor for stems, leaf photosynthesis, *V*_*cmax*_, *J*_*max*_, and *R*_*d*_ for leaves, and whole-plant conductance to water vapor for leaves and stems. All analyses and figures were made in R Studio v1.4.110 (R Studio) using R v4.0.3 (R Foundation for Statistical Computing).

## Results

### Photosynthetic response to diffuse light

Leaves differed in their photosynthetic rates in direct compared to diffuse light conditions. When considered relative to photosynthetic rates on the same leaf in direct light, there was a significant positive relationship between leaf photosynthetic rates and increasing diffuse light as induced by position of the light source on the integrating sphere (Figure 1a; *p* <0.00001, r^2^ = 0.48). For leaves, there is a clear increase of ∼ 1.5 μmol m^-2^ s^-1^ as light changes from predominantly direct to predominantly diffuse. Avocado stems perform low rates of photosynthesis whereby they re-assimilate respired CO_2_; there was no significant difference in net CO_2_ exchange rate under direct compared to diffuse light conditions (Figure 1b; t_23_ = 0.91, p = 0.37). When light respiration was subtracted from dark respiration to determine stem re-assimilation rates, there were also no differences between direct and diffuse light (Figure 1c; t_25_ = 0.92, p = 0.38).

**Figure 1.**
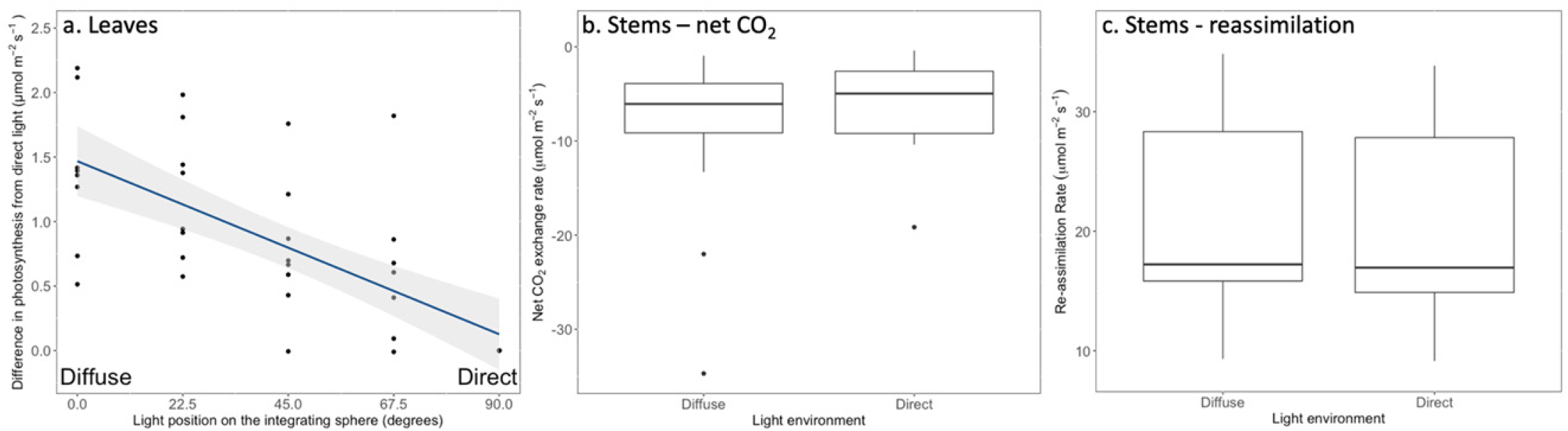
The effects of direct and diffuse light conditions on avocado leaf and stem gas exchange. The (a) difference in photosynthesis from direct light across different light positions on an integrating sphere. As light changes from direct to diffuse, leaves increase their photosynthetic rates. A single reference point exists for direct photosynthesis (a, bottom right) as all data was standardized against these data. Gray shading indicates the 95% confidence interval. The (b) stem net CO_2_ exchange and (c) re-assimilation rates in predominantly diffuse compared to predominantly direct light conditions. For stems, the net CO_2_ exchange and re-assimilation rates do not differ under diffuse or direct light conditions.

### Response of conductance to water vapor to diffuse light

For both leaves and stems, conductance to water vapor increased by ∼0.04 to 0.06 μmol m^-2^ s^-1^ in predominantly diffuse light conditions compared to predominantly direct light conditions (Figure 2). When considered relative to rates of conductance on the same leaf in direct light, there was a significant positive relationship between leaf conductance and increasing diffuse light as induced by position of the light source on the integrating sphere (Figure 2a; p <0.00001, r^2^ = 0.49). For stems, bark conductance to water vapor was significantly higher in diffuse compared to direct light conditions (Figure 2b; t_23_ = -2.77, p = 0.01).

**Figure 2.**
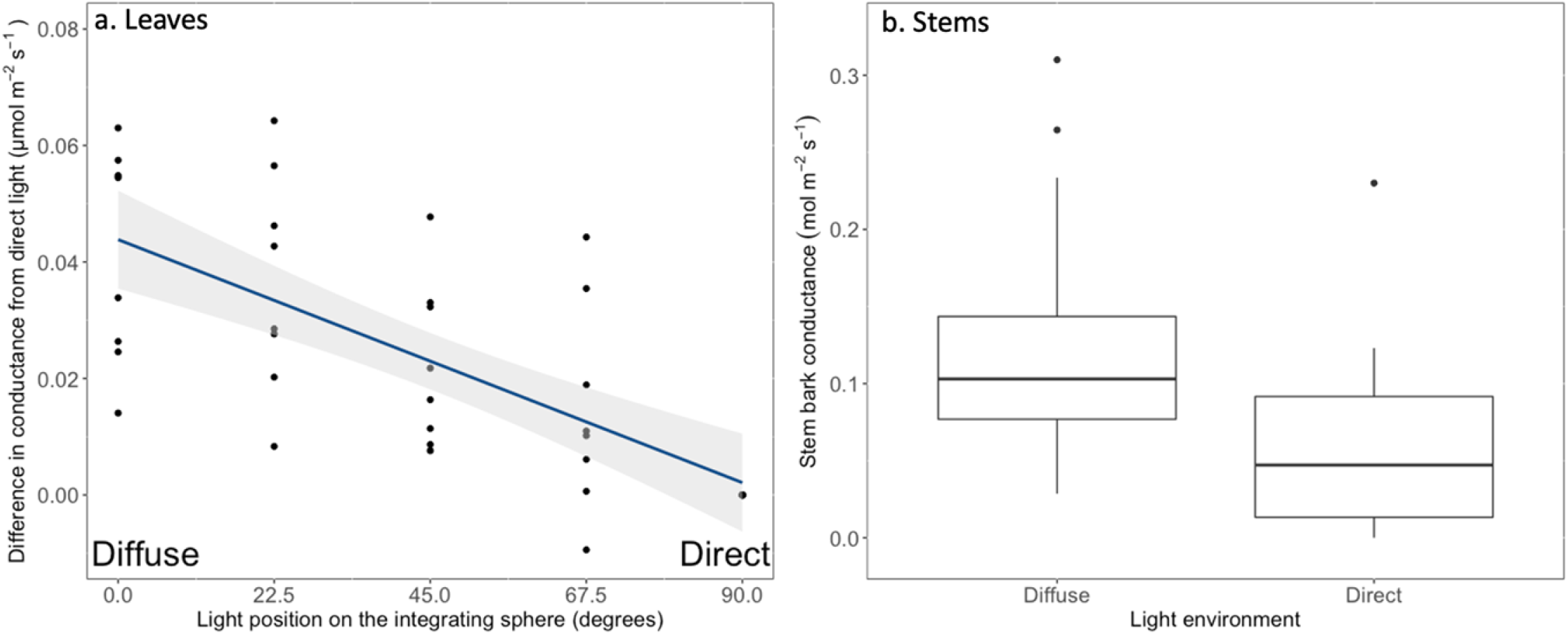
The effects of direct and diffuse light conditions on avocado leaf and stem conductance. The (a) difference between conductance in direct and diffuse light for paired measurements of individual leaves. As light changes from direct to diffuse, leaves increase their conductance. Gray shading indicates the 95% confidence interval. The (b) stem bark conductance in predominantly diffuse compared to predominantly direct light conditions. For stems, conductance is higher in diffuse light conditions

For leaves, the changes in stomatal conductance and photosynthesis under diffuse light conditions were closely related (Figure 3). The raw data of photosynthesis and conductance (Figure 3a) were fit using the square root model (p < 0.0001, r^2^ = 0.64). Because the data was collected on the same leaf under direct and diffuse conditions, we also paired the difference between diffuse and direct measurements per leaf and (Figure 3b) and fit a positive linear model (p < 0.0001, r^2^ = 0.62). For every increase in stomatal conductance of 0.02 μmol m^-2^ s^-1^ in diffuse light, there was a corresponding increase in photosynthesis of 0.53 μmol m^-2^ s^-1^.

**Figure 3:**
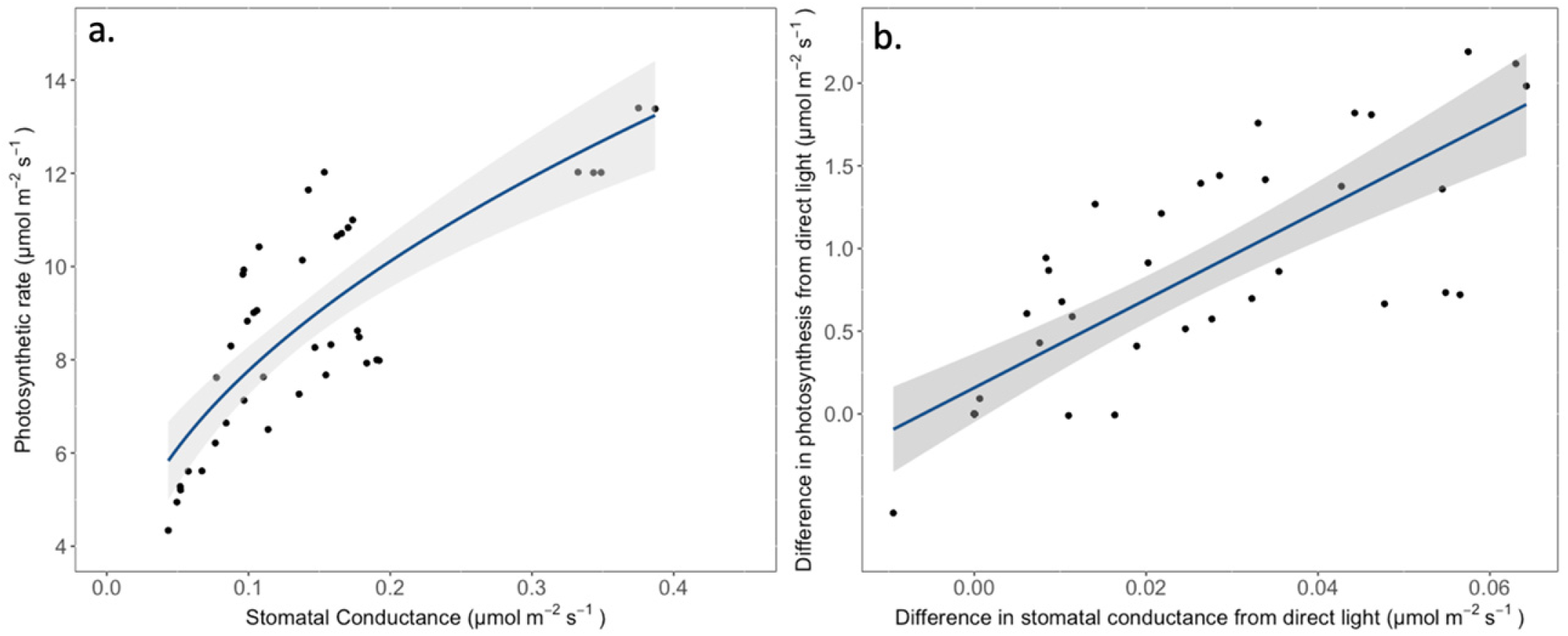
Relationship between photosynthesis and stomatal conductance with varying proportions of diffuse light. The changes to photosynthesis in direct or diffuse light appear to be driven by concurrent changes in stomatal conductance. This is observed in the relationships of (a) absolute values, or in (b) differences in gas exchange between direct and diffuse light measured on the same leaf. Gray shading indicates the 95% confidence interval.

### A-C_i_ curves

For leaves, we also explored the effects of diffuse light on leaf photosynthetic biochemistry. There were no significant differences between diffuse or direct light conditions on the maximum rate of Rubisco carboxylation (*V*_*cmax*_; t_29_= -0.38, p = 0.71), the maximum rate of electron transport (J_max_; t_29_= 0.41, p = 0.68), or dark respiration (R_d_; t_29_= 0.08, p = 0.94) (Table 1).

**Table 1.** Parameters of photosynthetic biochemical capacity extracted from photosynthetic response to CO_2_ curves. These parameters include the maximum rate of Rubisco carboxylation (*V*_*cmax*_), the maximum rate of electron transport (*J*_*max*_), and dark respiration (*R*_*d*_). Values represent means ± standard error.

| Light | $V_{cmax}$ ($\mu\text{mol m}^{-2} \text{s}^{-1}$) | $J_{max}$ ($\mu\text{mol m}^{-2} \text{s}^{-1}$) | $R_d$ ($\mu\text{mol m}^{-2} \text{s}^{-1}$) |
| --- | --- | --- | --- |
| Predominantly diffuse | $55.3 \pm 3.11$ | $95.2 \pm 5.92$ | $1.85 \pm 0.26$ |
| Predominantly direct | $56.6 \pm 2.35$ | $98.0 \pm 5.48$ | $1.83 \pm 0.20$ |

### Estimates of the contributions of leaves and stems to whole-plant carbon flux and conductance to water vapor

We measured leaf area, branch surface area, and trunk surface area (Table S1) and used these values to scale our leaf and stem level measurements to estimates of whole-plant conductance. Leaves accounted for 94.6 ± 0.48 % of conductance under direct light conditions, whereas stems accounted for the remaining 5.4 ± 0.48 % (Figure 4a,b). There were significant changes to the contribution of leaves and stems to whole plant conductance under diffuse light conditions. For conductance, stem fluxes contributed more to whole plant conductance under diffuse light conditions, increasing to 7.43 ± 0.65 % of total fluxes (t_11_ = 12.272, p < 0.0001).

**Figure 4.**
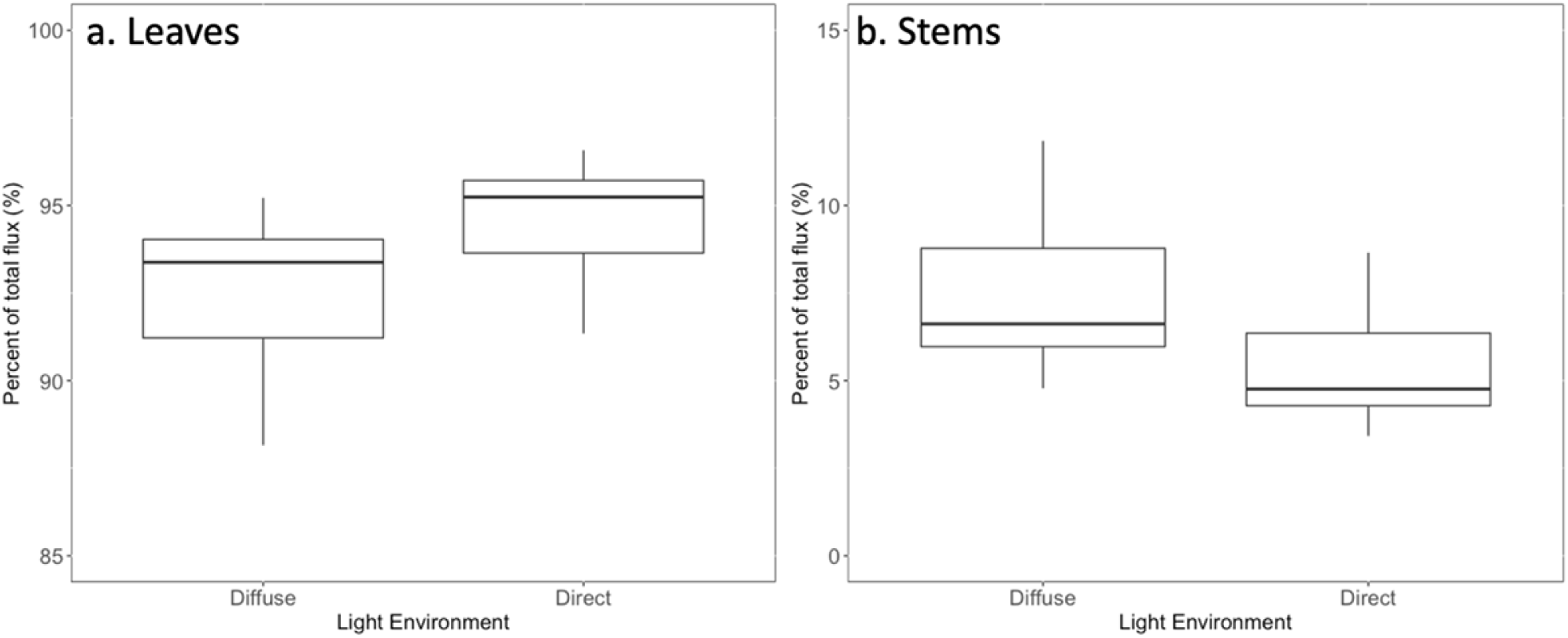
The percentage of whole-plant conductance coming from (a) leaves, or (b) stems under diffuse or direct light conditions. While stems are a relatively small contribution to whole plant conductance, the contribution is greater under diffuse light conditions.

## Discussion

We used an experimental manipulation to demonstrate that diffuse light leads to increases in leaf photosynthetic rates in avocado and that those changes are associated with stomatal opening, rather than photosynthetic biochemistry. Interestingly, we see a similar increase in stem bark conductance (but not photosynthetic rates) under diffuse light, despite stems lacking stomata. While others have suggested that reduced light transmittance under diffuse light should reduce photosynthetic rates (e.g., Earles et al. 2017), this is the first paper to demonstrate an explanation for why photosynthetic rates may *increase* under diffuse light. Finally, by scaling measurements to the entire plant, we demonstrate that stem bark conductance contributes proportionally more to whole-plant conductance under diffuse light conditions. Below, we explore potential explanations and implications for these findings.

### Think you might need a section here discussing variation of photosynthesis with intermediate (varying) amounts of light, so that you can parallel the questions you set up in the introduction

#### Why is leaf conductance changing under diffuse light?

By manipulating the proportion of diffuse light reaching the leaf surface while holding the quantity of light constant, we were able to observe a relationship between leaf conductance and photosynthetic rates. It is well established that stomata respond to light, particularly blue and (to a lesser extent) red wavelengths (Shimazaki et al. 2007), despite the majority of this light being absorbed in upper mesophyll layers. Blue light is sensed in guard cells by phototropins and can be triggered at very low levels (as low as 10 μmol m^-2^ s^-1^; Kinoshita et al. 2001). If the major light sensing phototropins are in guard cells on the abaxial surfaces of leaves, then reduced light transmittance in diffuse light should lead to reduced conductance. Of course, if the change in light transmittance is minimal, then the guard cell phototropins could still detect some blue light leading to no change in stomatal conductance. Neither mechanism could explain why in this study, and with our previous work (Berry et al. 2018, Ellertson et al. 2022), we observe that diffuse light often increases photosynthetic rates and conductance to water vapor under diffuse light. Therefore, the changes to conductance are likely coming from cues other than direct stimulation by light.

Guard cell opening with red light is largely considered a minor process but, interestingly, is not dependent on light reaching guard cells (Matthews et al. 2020). As red light stimulates mesophyll photosynthesis, sugars can be transported through H+/sucrose symporters directly into guard cells to induce opening (Ando et al. 2018). Additionally, guard cells react to leaf intercellular CO_2_ (*c*_*i*_) so that increased photosynthetic activity under red light would further promote guard cell opening through reduced *c*_*i*_ (Roelfsema et al. 2002). This hypothesis could explain our results. Leaves were exposed to red and blue light and therefore mesophyll photosynthesis was still very active during diffuse light conditions. But, for conductance to be greater under diffuse light would require conditions that increase photosynthesis during diffuse light. For that we need to assess other environmental considerations that might be driving this response.

Of course, it is well established that photosynthetic rates and stomatal conductance respond to cues other than light, such as temperature, water supply and atmospheric water demand. Even though our gas exchange measurements held leaf temperature and humidity constant, it is plausible that subtle changes to these parameters may aid in explaining diffuse light carbon fluxes. Broadly, higher leaf temperatures lead to increased stomatal conductance, presumably to increase available CO_2_ and evaporative cooling (Schulze et al. 1973; Lu et al. 2000; Ameye et al. 2012, Urban et al. 2017). Other studies have shown that diffuse light from greenhouse coatings reduce air and leaf temperatures, which would suggest stomatal closure, not the opening we see here (Dueck et al. 2012, Ellertson et al. 2022). However, slight decreases in leaf temperature could decrease vapor pressure deficit in the leaf boundary layer, which is known to increase stomatal conductance (McAdam et al. 2015, Pantin et al. 2018). This combined temperature and vapor pressure deficit response could interact to induce increased stomatal conductance under diffuse light conditions.

#### Why would stems have higher bark conductance with no changes to net CO_2_ exchange?

In leaves, the increased conductance led to greater carbon gain. In stems, we observed increased conductance and no concomitant change in carbon gain. There are several reasons why we might not expect a tight coupling of stem bark conductance to water vapor and photosynthesis under diffuse light. First, because avocado stems perform recycling photosynthesis (carbon assimilation is utilizing CO_2_ released from respiration Nilsen 1995; Ávila-Lovera et al. 2014), carbon assimilation is not relying on the diffusion of atmospheric CO_2_ through the bark for photosynthesis. Second, avocado stems do not have stomata and therefore conductance reflects stem periderm permeability that is governed exclusively by the vapor pressure gradient between the stem and the atmosphere. There is no evidence that the vapor pressure gradient significantly differed between the direct and diffuse light treatments in which we made our experimental measurements of conductance were made. And finally, all light may, functionally, be diffuse for chlorophyll in photosynthetic stems. As stems have multiple protective layers of cellular tissue that light must pass through, direct light hitting the surface may be diffuse by the time it reaches chlorophyll. If direct and diffuse light transmit similarly through stems, then the same amount of energy would be supplied to chloroplasts. While the mechanism that would explain the change in stem bark conductance in diffuse light is unclear, the lack of stomata allow for the decoupling of stem conductance and photosynthesis.

#### Whole-plant conductance - tradeoffs in leaf and stem conductance

Green stems can play an important role in whole-plant carbon balance (Nilsen 1995, Ávila-Lovera et al. 2014), particularly in seasonally dry systems where plants lose their leaves or do not have leaves for prolonged periods of drought (Ávila-Lovera et al 2019). Avocado differs from these species because leaves turnover, but generally do so simultaneous to the flushing of new leaves. Thus, stems are never the sole source of assimilated carbon. In our one-year-old avocado trees, stems accounted for 5.4 ± 0.48 % of whole plant conductance under direct light conditions (Fig. 4). The role of stems to whole plant conductance increased under diffuse light to 7.43 ± 0.65 %. Even though this percent may seem small, stem bark conductance to water vapor is associated with drought performance, with greater values of conductance inducing greater mortality of tropical tree seedlings (Wolfe 2020). This increase in importance of stems may also lead to significant gains in carbon over the course of a growing season. Thus, small changes in whole-plant carbon and water fluxes driven by stems could have major impacts on plant survival.

While photosynthesis and conductance remained tightly coupled for leaves with varying proportions of diffuse light, there was a clear tradeoff as increasing proportions of diffuse light were significantly related to decreases in water-use efficiency (Fig. S1). However, stems do not have this issue, as carbon availability is decoupled from water loss. Green stems of desert plants and plants from the dry tropics have been observed to be more water-use efficient than leaves (Ehleringer et al 1987, Nilsen and Sharifi 1997, Smith and Osmond 1987, Ávila and Tezara 2018). However, it has been recognized that conductance and transpiration rates can be high in stems of deep-rooted plants (Gibson 1996, 1998). In this study, increased conductance may be acting to regulate temperature. This leads to a tradeoff between reduced leaf temperature and increased water loss.

#### Mechanisms of diffuse light photosynthesis in response to projected changes in climate

Whether diffuse light is due to aerosols, pollution, clouds, forests canopies, or greenhouse materials, it is a ubiquitous part of the plant experience. Globally, the proportion of diffuse light is increasing (Mercado et al. 2009), leading to important questions about what drives changes to plant function under diffuse light and the environmental implications of this response. Our recent work has sought to isolate the effects of diffuse light on plant function to demonstrate changes that can both increase and decrease productivity.

The current study begins to address multiple questions about the next phase of this work. Why would leaf photosynthesis increase under diffuse light? Here, we find that conductance plays a key role; however, the generality of this response across species remains to be established. How does the response to diffuse light vary across plant organs? We find that green stem and leaf conductance respond similarly to diffuse light without the associated effects on carbon balance for green stems. How does diffuse light integrate to drive whole plant conductance? We find that green stem conductance increased in importance to whole plant conductance under diffuse light conditions. Our results suggest compelling questions about the interactive effects of diffuse light with other environmental variables, particularly atmospheric humidity. While our approach has attempted to hold these variables constant, in natural conditions these variables may be influential to carbon-water tradeoffs. At broad scales, diffuse light reduces temperatures and vapor pressure deficits (Gu et al. 2002, Alton et al. 2007), which should increase conductance. Thus, the extent to which the combination of responses to temperature, vapor pressure deficit, and diffuse light affect conductance and carbon-water tradeoffs in natural conditions requires further research to resolve.

Our work has demonstrated that diffuse light is a fundamental environmental variable that affects plant function. As diffuse light broadly increases, some species may increase photosynthesis through increased conductance, but thus experience reduced water-use efficiency. The extent to which the functional response to diffuse light is mediated by decreased soil moisture (i.e., drought), increased global temperatures, or increased vapor pressure deficit is unknown, but could be crucial in unpacking the broader implications. We now need to integrate these responses into our complete understanding of plant responses to climate change.

## Acknowledgements

We want to thank the support staff that allowed us to conduct this research project during the height of the Covid-19 pandemic. This project was funded by USDA-NIFA Award# 2020-67014-30916 to ZCB and GRG and USDA-NIFA Award# 2020-67014-30915 to EAL and GRG.

## Author Contributions

ZCB, EAL, and GRG designed the research, KE, ZCB, and EAL performed the work and analyzed the data. All authors contributed to writing and editing of the manuscript.

## Supplemental Information

**Figure S1:**
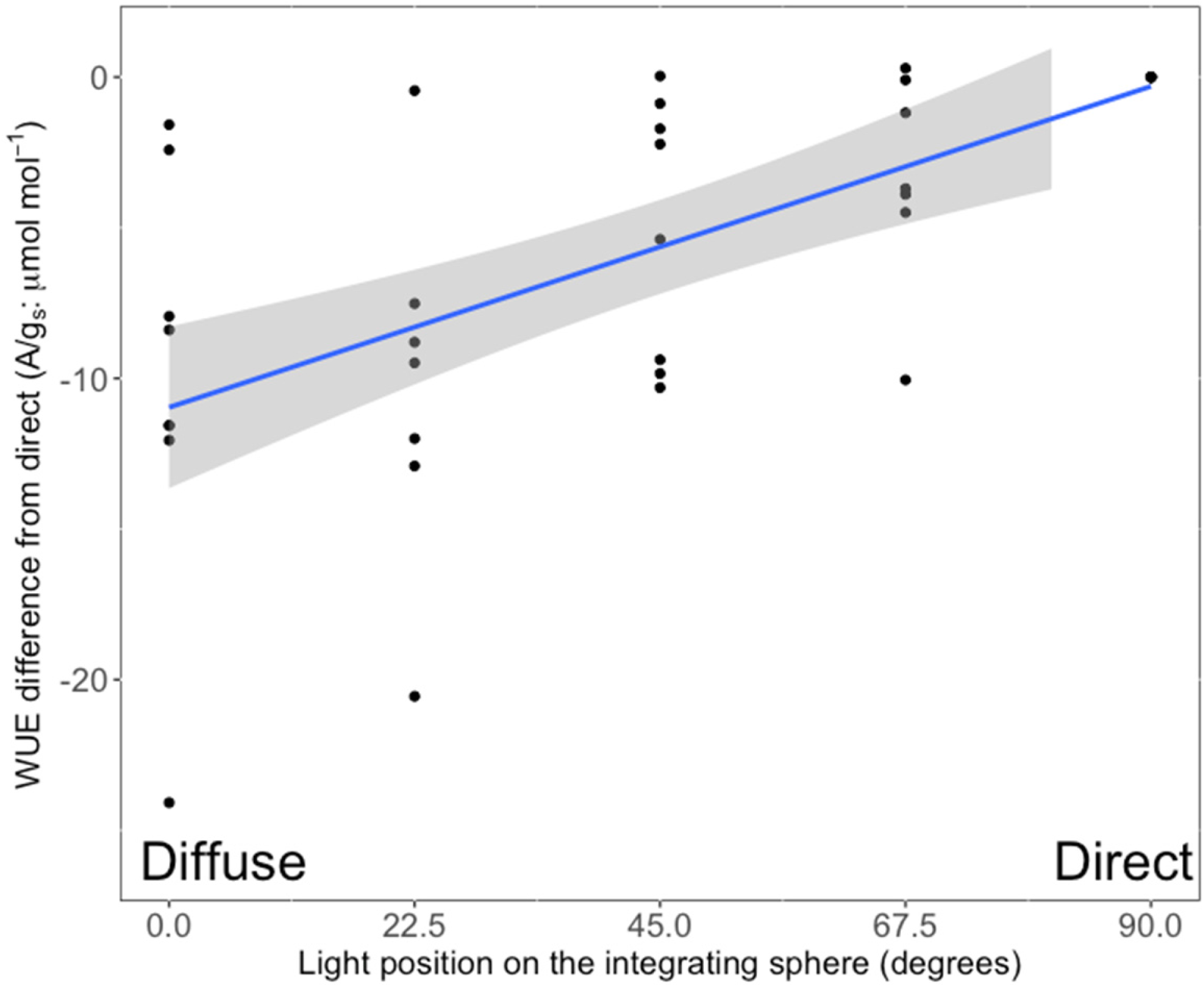
The change to leaf intrinsic water-use efficiency (iWUE) as light changes from predominantly direct to predominantly diffuse. Because of the increases in stomatal conductance (i.e. Figures 2 & 3) under diffuse light conditions, the iWUE decreases.

**Table S1.** Total plant leaf area, branch surface area, and proportion of leaf to branch area ratio used for scaling leaf and branch gas exchange to whole-plant conductance.

| Individual | Total plant leaf area (m<sup>2</sup>) | Branch surface area (m<sup>2</sup>) | Leaf area/branch area ratio |
| --- | --- | --- | --- |
| 1 | 33.89 | 3.82 | 8.85 |
| 2 | 22.05 | 3.30 | 6.68 |
| 3 | 19.60 | 3.77 | 5.21 |
| 4 | 43.06 | 3.50 | 12.31 |
| 5 | 45.45 | 3.97 | 11.44 |
| 6 | 41.94 | 4.25 | 9.87 |
| 7 | 32.28 | 4.06 | 7.96 |
| 8 | 42.99 | 4.05 | 10.63 |
| 10 | 18.42 | 2.84 | 6.49 |
| 11 | 66.73 | 4.79 | 13.92 |
| 12 | 34.36 | 3.48 | 9.87 |

